# AIDeveloper: deep learning image classification in life science and beyond

**DOI:** 10.1101/2020.03.03.975250

**Authors:** Martin Kräter, Shada Abuhattum, Despina Soteriou, Angela Jacobi, Thomas Krüger, Jochen Guck, Maik Herbig

## Abstract

Publications on artificial intelligence (AI)-based image analysis have increased drastically in recent years. However, all applications use individual solutions highly specialized for a particular task. Here, we present an easy-to-use, adaptable, open source software, called AIDeveloper (AID) to train neural nets (NN) for image classification without the need for programming. The software provides a variety of NN-architectures that can be simply selected for training. AID allows the user to apply trained models on new data, obtain metrics for classification performance, and export final models to different formats. The working principles of AID are first illustrated by training a convolutional neural net (CNN) on a large dataset consisting of images of different objects (CIFAR-10). We further explore the potential of AID by training a model to distinguish areas of differentiated and non-differentiated mesenchymal stem cells (MSCs) in culture. Additionally, we compare a conventional clinical whole blood cell count with a whole blood cell count performed by an NN-trained, using a dataset of more than 1.2 million images obtained by real-time deformability cytometry, delivering comparable results. Finally, we demonstrate how AID can be used for label-free classification of B- and T-cells derived from human blood, which currently requires costly and time-consuming sample preparation. Thus, AID can empower anyone to develop, train, and apply NNs for image classification. Moreover, models can be generated by non-programmers, exported, and used on different devices, which allows for an interdisciplinary use.

## Introduction

Since the development of the first microscope, progress in life science has become dependent on image acquisition and processing. Over the years, extensive research has led to the development of tools used for quantitative analysis of information in microscopic images. Software such as Cellprofiler (Jones et al., 2008), Fiji (Schindelin et al., 2012) and ImageJ (Schneider et al., 2012), which are widely distributed and used among scientists, allow the user to easily process and quantify features from images. However, quantification using these tools is constrained to a set of predefined features that often limit the extent of information that can be extracted from the images. In recent years, the emergence of machine learning (ML) methods, such as deep learning (DL) which uses neural nets (NN), substantially augmented the scope of image processing, quantification, segmentation, and classification. The main advantage of the DL approach is that it does not rely on handcrafted, predefined features, but rather automatically finds a set of optimal features. This can be especially helpful for complex image classification tasks where relevant features are not obvious to the human eye.

Recent publications demonstrated the applicability of DL for image-based object identification in complex biological samples. For example, thrombocyte clusters were identified in human blood samples (Nitta et al., 2018), cell lineage differentiation was predicted during hematopoietic stem cell development (Buggenthin et al., 2017), skin cancer was classified on dermatologist-level (Esteva et al., 2017), and mitotic cells were detected in histology images (Cireşan et al., 2013; Giusti et al., 2014). The latter showed DL to even outcompete histologists in terms of accuracy (accuracy = number of correctly classified images per total number of images). However, only customized, task-specific algorithms are currently available, as the accessibility and utilization of DL algorithms requires distinct programming skills. Thus, there is an increasing demand to make DL-based image processing accessible for the general user.

Here, we present AID (artificial intelligence developer), a flexible ready-to-use software platform to train, evaluate, and utilize NNs for image classification problems. AID covers the entire work flow of image processing and analysis: from the assembly of datasets and the optimization of NN parameters, to the application of the generated NN to unclassified image sets. A simple user interface allows the user to load different image formats and to visually assess them before and after image size equalization. For training, the user can either choose NNs of different complexities or use custom-built NNs. In addition, the interface allows the use of pre-trained models to transfer the learning process, i.e., when insufficient training data is available (Tan et al., 2018) or to shorten the training step.

To demonstrate the software’s potential, we trained a convolutional neural net (CNN) on CIFAR-10, a dataset containing a collection of images from 10 different classes (Krizhevsky, 2009). We reached a testing accuracy as high as 88% on RGB and 83% on grayscale images. Furthermore, we demonstrate the application of AID for broad biomedical research. First, we trained a model to detect differentiated adipocytes using a relatively small dataset containing only 46 labelled bright-field microscope images. Next, we show the utility of AID for very large datasets by using 1.2 million images of blood cells obtained with real-time deformability cytometry (RT-DC) (Otto et al., 2015), in order to generate an automatic whole blood cell count. We trained a model to recognize thrombocytes, lymphocytes, red blood cells, monocytes, neutrophils, and eosinophils based on bright-field images acquired using RT-DC. We validated the model by comparing the result to a conventional whole blood count generated by a technique frequently used in clinical practice, which agreed well. Finally, we demonstrate that the tools provided by AID master even challenging classification tasks by training a classifier to distinguish B- and T-cells based on bright-field images from RT-DC. The resulting model reaches a classification performance that is state-of-the-art for label-free approaches (Nassar et al., 2019; Yoon et al., 2018, 2017). With the demonstrated utility for a wide range of potential image analysis tasks, AID is a ready-to-use software package for anyone who wants to start exploring the power of AI-based image analysis for their own research without the need for any programming skills.

## Results

### Using AID to classify natural images from CIFAR-10 and Fashion-MNIST

AID enables anyone to apply DL for image classification as it guides the user through the entire project pipeline, starting from loading and assembling a dataset, proceeding with training and evaluating a DL model, and ending with classifying new sets of images (Figure 1A, Figure 1— figure supplement 1, Video 1). Images for validation and training are simply dragged and dropped into a designated area of the user interface where they are converted to a uniform data format. The validation set is used after every training iteration to validate the generated model. A library of seven different multilayer perceptrons (MLPs) and 23 convolutional neural nets (CNNs) of a wide range of complexity is available for choosing a model architecture (Figure 1—figure supplement 2). In addition, custom-built CNNs as well as pre-trained models are supported. Pre-trained models can be used either for classifying new sets of images or for re-purposed training on a different classification task, a technique termed transfer learning (Tan et al., 2018). At the initiation of a training process, AID automatically generates the neural net architecture according to the requested input and output dimensions. Image sizes are adjusted either by cropping or padding. AID offers the adjustment of a range of training parameters before and during the training process. These parameters are known as hyper-parameters and include different image augmentation options, learning rate, and dropout rate (Video 1). Example images can be visualized before and during the training process in order to assist the adjustment of image augmentation parameters (Video 1). The accuracy and the validation accuracy are plotted in real-time after each training iteration. Furthermore, F1 score, precision, recall, support, receiver operating characteristic curve, precision-recall curve, and further common metrics are shown and can be exported (Swets, 1996). Once a suitable model is obtained (i.e., according to validation accuracy) it can be loaded into AID to assess its performance on testing data (Video 1). Since developers might want to use a trained model in a different framework, AID also provides conversion tools to protocol buffer format (TensorFlow) (Abadi et al., 2016) and ONNX (https://github.com/onnx/onnx) (Video 1). All analyses for this work are performed using AID on a standard consumer PC.

**Figure 1:**
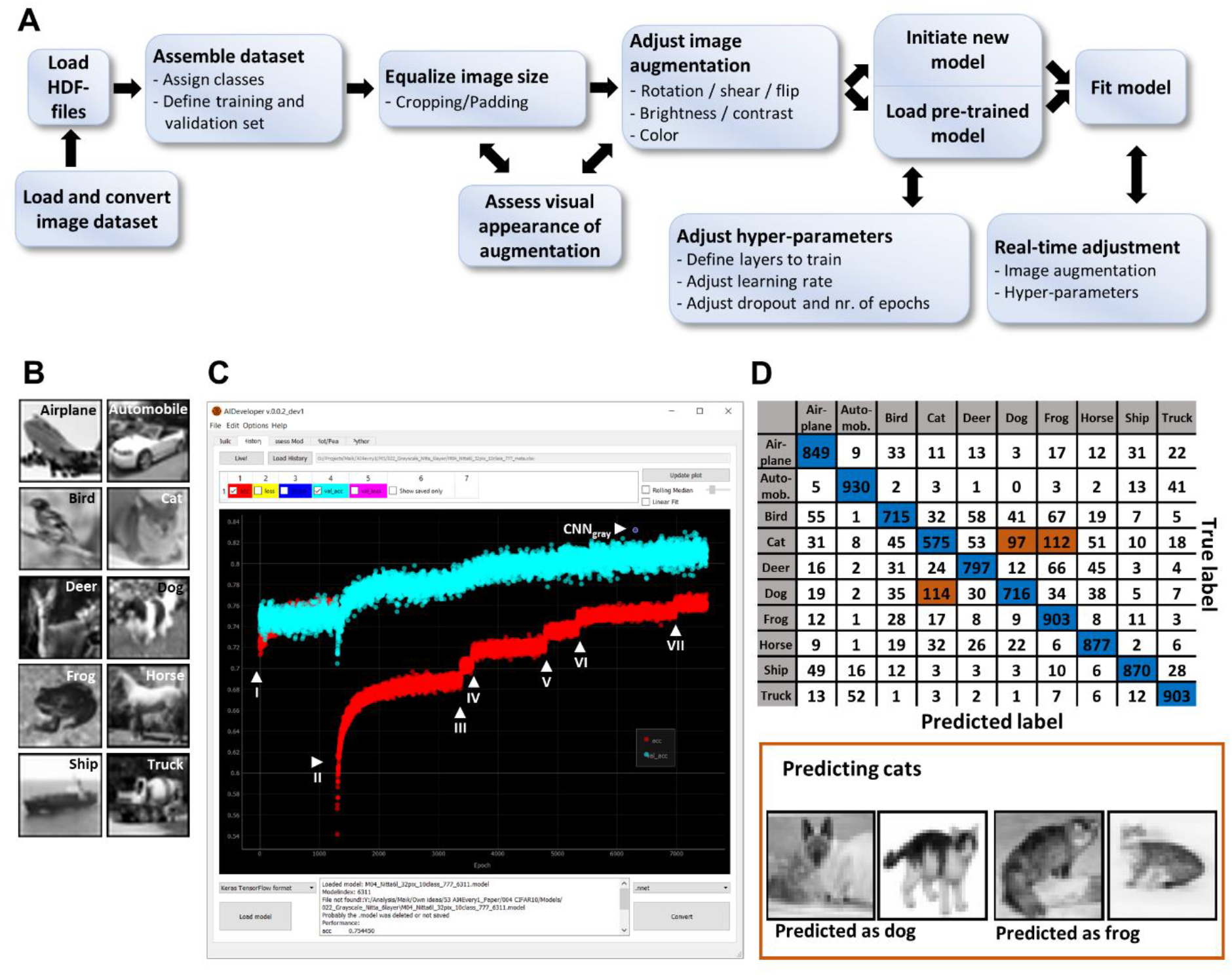
AIDeveloper user interface and workflow. (**A**) A representative workflow of setting up a training process. (**B**) Representative grayscale images of all CIFAR-10 classes (out of 6000 images per class). (**C**) Screenshot showing the “history”-tab of AID, which was used to load the training history file of the training process for grayscale images. The scatterplot shows the accuracy (red dots) and the validation accuracy (cyan dots) for each training iteration (also called ‘epoch’). Arrowheads indicate seven different real-time user adjustments (I to VII) of image augmentation or hyper-parameters. CNN_gray_ indicates the model at epoch 6311, which reaches the maximum validation accuracy. (**D**) A confusion matrix indicating the true and the predicted label when classifying the testing set of CIFAR-10 using CNN_gray_. Matrix items with blue and orange color indicate correctly and incorrectly predicted classes, respectively. Representative images of incorrect predictions from model CNN_gray_ on CIFAR-10 of class “cat” is shown. The testing accuracy is 81.2%.

To illustrate the software’s potential, CIFAR-10, a common dataset for benchmarking new image classification algorithms, was used. CIFAR-10 contains 6000 different images of each of the following classes: airplane, automobile, bird, cat, deer, dog, frog, horse, ship, and truck (Figure 1B). 4800 images were used as training set, 200 images of each class as validation set and 1000 images as testing set. We chose a CNN architecture with 4 convolutional layers (Figure 1—figure supplement 2N) (Nitta et al., 2018) and used grayscale images to reduce the computational time and accelerate the training process. During training, the image augmentation parameters were adjusted seven times. This caused immediate changes in both the accuracy and the validation accuracy. Importantly, training proceeded with an overall improvement of the validation accuracy, and the best model reached a validation accuracy of 83.2% (indicated as CNN_gray_ in Figure 1 C). Furthermore, we trained the same CNN-architecture using the original RGB images, resulting in a model with a validation accuracy of 87.9% (CNN_RGB_). Both models (CNN_gray_ and CNN_RGB_) were then applied to the testing set. We obtained a testing accuracy of 81.2% and 84.7% for CNN_gray_ and CNN_RGB_, respectively. The resulting confusion matrix for CNN_gray_ indicates that the distinction between animals, especially classification of cats, dogs, and frogs, was the most erroneous (highlighted in orange in Figure 1D). In AID, confusion matrices are interactive and allow the user to visualize the respective images for each matrix position (Figure 1D bottom panel).

To further evaluate AID’s performance, we resorted to Fashion-MNIST (Xiao et al., 2017) an image dataset of 10 different classes of fashion items. The dataset contains 7000 images for each class and we used 5800 images for training, 200 for validation and 1000 for testing. The CNN_gray_, previously trained on CIFAR-10 images (32 x 32 pixels) was further trained on Fashion-MNIST images (28 x 28 pixel). To allow transfer learning, we utilized the image scaling option in AID to match image sizes. Initially, only the last layer of the pretrained CNN was left trainable while all other layers were frozen. By gradually unfreezing all layers (Yosinski et al., 2014) during training, we reached a robust classification model with an accuracy of 95.1% on the validation set and 93.8% on the testing set. The training progress and a resulting confusion matrix is visualized in Fig. 1 – figure supplement 3.

### AID in life science and its potential for clinical diagnostics

AID’s ability to train models to natural images could be applied in mobile app development or autonomous driving, where large numbers of natural (everyday-life) images are encountered (Caesar et al., 2019). Here, we focus on demonstrating its potential for the classification of images in life science and clinical diagnostics that also encounter the challenges of processing large image datasets.

Mesenchymal stem cells (MSCs) hold a great potential for the future of cell-based therapeutic approaches. However, prior to the transplantation it is essential to characterize MSCs and assess their differentiation potential. One classical approach includes MSC differentiation into adipocytes, followed by histological analysis with the lipid dye Oil Red O and manual quantification of the stained cells (Majumdar et al., 1998; Oswald et al., 2004). Here, we acquired bright-field images (320 x 320 pixels) of Oil Red O-stained adipocytes from different positions in the cell culture well (Figure 2A, B). Since brightness, color, and distribution of Oil Red O staining varied significantly (Figure 2B and Figure 2—figure supplement 1), a trained person was asked to visually grade and mark the differentiated areas of the acquired images. We then used the information from the manual labelling and masked the differentiated areas with a uniform green color (RGB: 0, 255, 0) (Figure 2C). Each acquired image was divided into 100 tiles of 32 x 32 pixels and used as a training dataset (Figure 2C). Tiles with less than 5 labelled pixels were assigned to class 0 (“without differentiation”) while all other tiles are assigned to class 1 (“with differentiation”).

**Figure 2:**
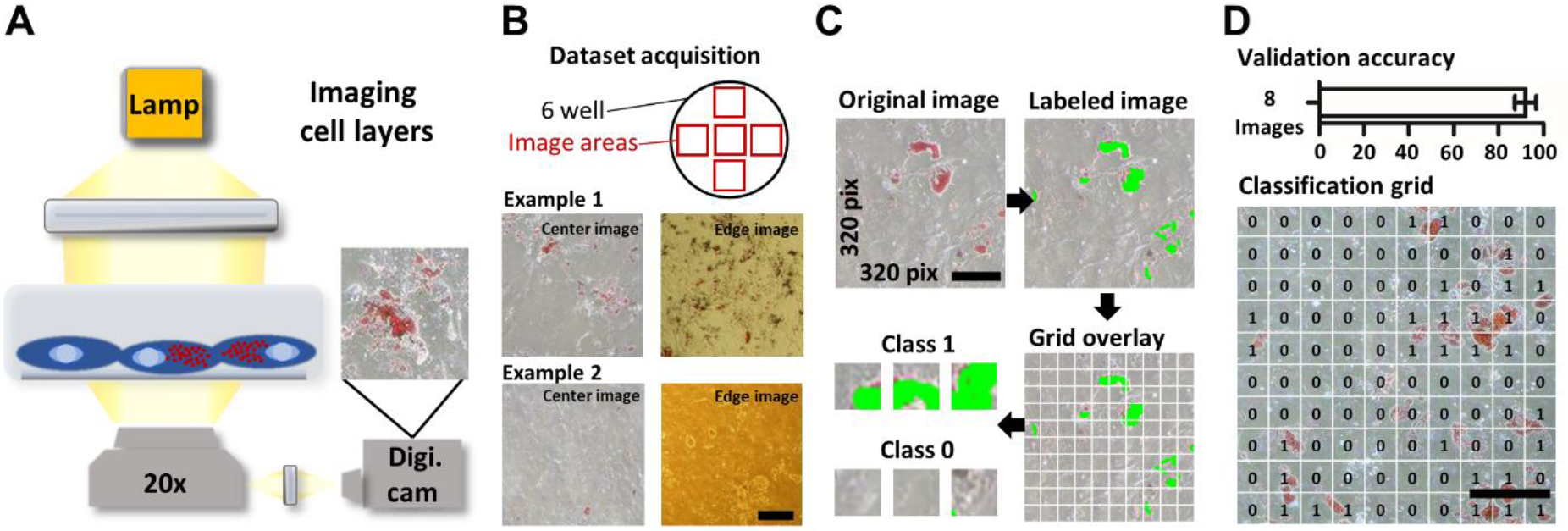
Classifying image tiles containing adipogenic differentiated mesenchymal stromal cells. (**A**) Schematic representation of a light microscope to image cells in 2D culture. Human mesenchymal stromal cells were induced to differentiate into the adipogenic lineage and imaged following Oil Red O staining. (**B**) Image acquisition strategy. Cells were cultured in a six-well plate and 5 fixed positions within each well were imaged. Representative images of the center and edge position of two different examples are shown, indicating the variability in image color and staining quality. (**C**) Image-processing pipeline to obtain training data. Areas of cell differentiation were labeled and the original 320 x 320 pixels images were divided into 100 tiles (32 x32 pixels). Tiles containing more than 5 labeled pixels were assigned to class 1, others to class 0. (**D**) The bar graph presents the averaged validation accuracy over 8 images ± S.D. The image presents the classification of a new image neither contained in the trainingset nor the validation set. The numbers indicate whether a tile was predicted to belong to class 1 (“with differentiation”) or class 0 (“without differentiation”). Scale bars = 50 μm.

For the classification task, we designed a CNN with 6 convolutional layers, 4 fully connected layers, and residual connections between layers (Figure 1—figure supplement 2M). A transfer learning approach was chosen, where the model was first trained on RGB images from CIFAR-10 and subsequently optimized for the task of distinguishing tiles with and without differentiation. The final model was validated using eight images, resulting in a validation accuracy of 92.1% (Figure 2D and Figure 2—figure supplement 1). Finally, we tested the model using an unlabeled image not contained in the training-nor validation set. As shown in Figure 2D the tiles classified to class 1 (“with differentiation”) were in good agreement with stained regions.

High-throughput imaging allows fast screening of large amounts of biological samples. These images potentially include information that is useful for clinical diagnostics, creating a need for fast and automated image classification. AI-based image classification is a highly useful technique for advancing the field of image-based real-time diagnosis. Here, we combined real-time deformability cytometry (RT-DC) and AID to generate a label-free, image-based blood cell count. RT-DC is an imaging flow cytometer where bright-field images of cells in flow are captured by a high-speed camera (Figure 3A) at rates of 100 – 1,000 cells/s. In order to increase the frequency of leucocytes, whole blood was first depleted from RBCs by dextran-sedimentation (Bøyum, 2008; Quach and Ferrante, 2017). As recently published by Toepfner et al., the individual cell populations can be identified simply by considering cross-sectional area and brightness, calculated as the average grayscale value of all pixels belonging to the cell (Figure 3B). These prameters are sufficient to distinguish thrombocytes, RBCs, RBC doublets, lymphocytes, monocytes, neutrophils, and eosinophils (Toepfner et al., 2018). To avoid incorrect labelling, a conservative gating was applied, by excluding events where a distinction was not obvious. Basophils are excluded from our model as they are difficult to distinguish based only on area and brightness. Furthermore, they are very rare and we were not able to label a sufficient number of cells for training. We assembled a training and a validation set containing approximately 1.2 million images and trained a LeNet5 (Lecun et al., 1998) architecture (Figure 1—figure supplement 2D), reaching a validation accuracy of 97.3%. Testing data was acquired by measuring 17 additional blood samples using RT-DC and comparing to a conventional blood count measured in parallel under clinical settings. The trained model was applied to classify the images from the RT-DC experiments. The resulting cell count was comparable to the conventional blood count (Figure 3C). Using RT-DC, an additional population of RBC doublets was found, which is not reflected in the conventional blood count.

**Figure 3:**
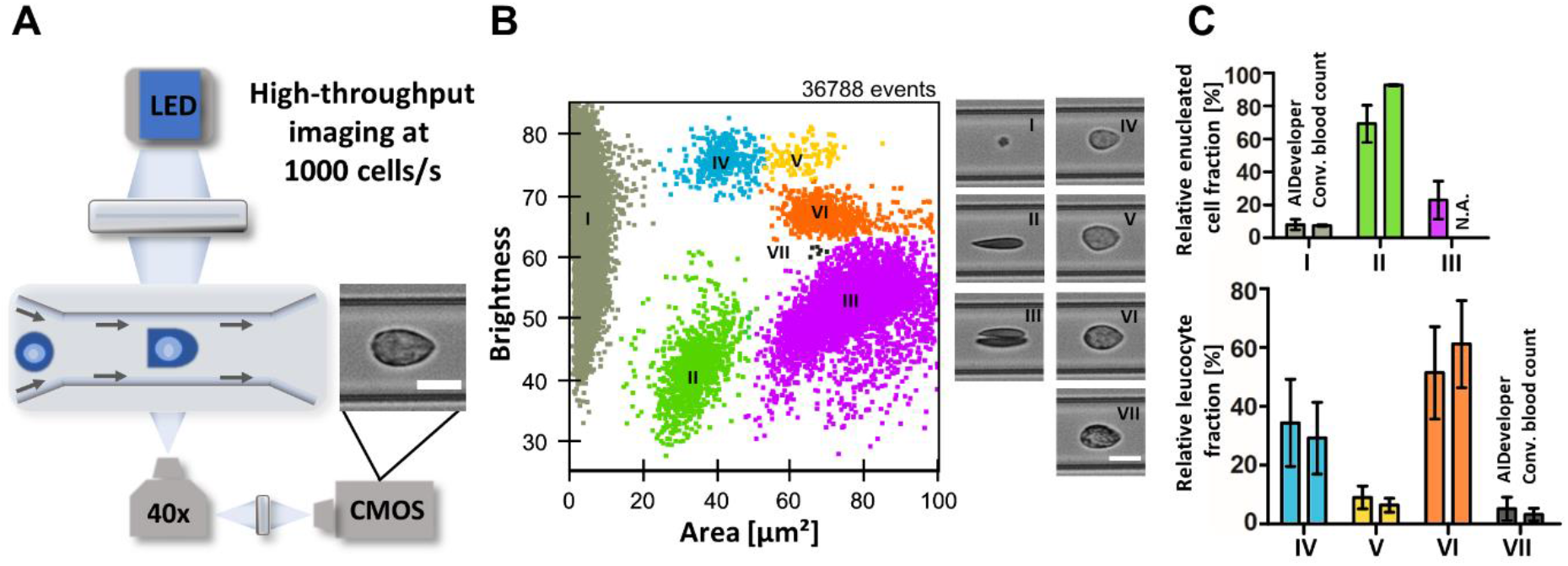
Image-based whole blood count using RT-DC and AID. (**A**) Schematic representation of RT-DC, a high-throughput imaging technology. A cell suspension is flushed through a channel constriction in a microfluidic chip. Cells are illuminated by an LED and recorded by a high-speed camera. Multiple parameters including area and average brightness of the cells are determined in real-time. Scale bar = 10 μm (**B**) The brightness versus area scatter-plot of whole blood measurements is used to distinguish populations of the major blood cells (I thrombocytes, II erythrocytes, III erythrocyte doublets, IV lymphocytes, V monocytes, VI neutrophils, and VII eosinophils) (Toepfner et al., 2018). Corresponding images of each population highlight the phenotype of these cells. Manual gating of these populations was carried out to assemble a dataset for training a CNN to perform an image-based whole blood count. Scale bar = 10 μm (**C**) The bar-graphs present the relative fraction of enucleated cells (I thrombocytes, II erythrocytes, and III erythrocyte doublets) as well as the leucocytes (IV lymphocytes, V monocytes, VI neutrophils, and VII eosinophils), determined using the CNN and a conventional blood count. Mean ± S.D. of 17 independent blood measurements is displayed.

To emphasize the potential of DL and AID in the context of blood analysis, we set out to classify lymphocytes into B- and T-cells label-free, a task not yet feasible routinely in clinical diagnostics. To discriminate between B- and T-cells in whole blood, we used real-time fluorescence and deformability cytometry (RT-FDC) (Rosendahl et al., 2018). This technique is similar to RT-DC, but allows for fluorescence detection in parallel to imaging. We generated a labelled dataset from 3 healthy donors, by using a panel of three fluorescent antibodies (ABs) specific for each cell type; “cluster of differentiation” 19 (CD19) for B-cells; CD3 for T-cells and CD56 for NK-cells, a subset of T-cells. As previously described, we identified lymphocytes based on area and brightness (blue in Figure 4A). B- and T-cells were detected based on expression of the different fluorescent markers as shown in Figure 4A. Training and validation sets were assembled using data from three donors. We used a 4^th^ measurement from a different healthy donor as testing dataset. Acquired images for B-cells (CD19^+^ events) and T-cells (CD19^+^ or CD56^+^ events) were loaded into AID and assigned to individual classes. CNN_gray_ (Figure 1C and Figure 1—figure supplement 2N), was used to apply transfer learning. The parameters of the first convolutional layer were set constant and training of the rest of the model-parameters was continued using image data of B- and T-cells (Figure 4B). The final model reached a validation accuracy of 89.3%. We used AID to apply the model on testing data and obtained the following scores: testing accuracy = 86.2%, F1 score = 89.3%, precision = 92.3%, recall = 86.5% and an area under curve (AUC) of the receiver-operating characteristic (ROC) and the precision recall curve of 94% and 97%, respectively. The probability histogram of the testing-set (Figure 4C) indicates that above a threshold of p_T_ = 0.9, 98.3% of the events are classified as T-cells and below p_T_ = 0.1, 95.9% of the events as B-cells.

**Figure 4:**
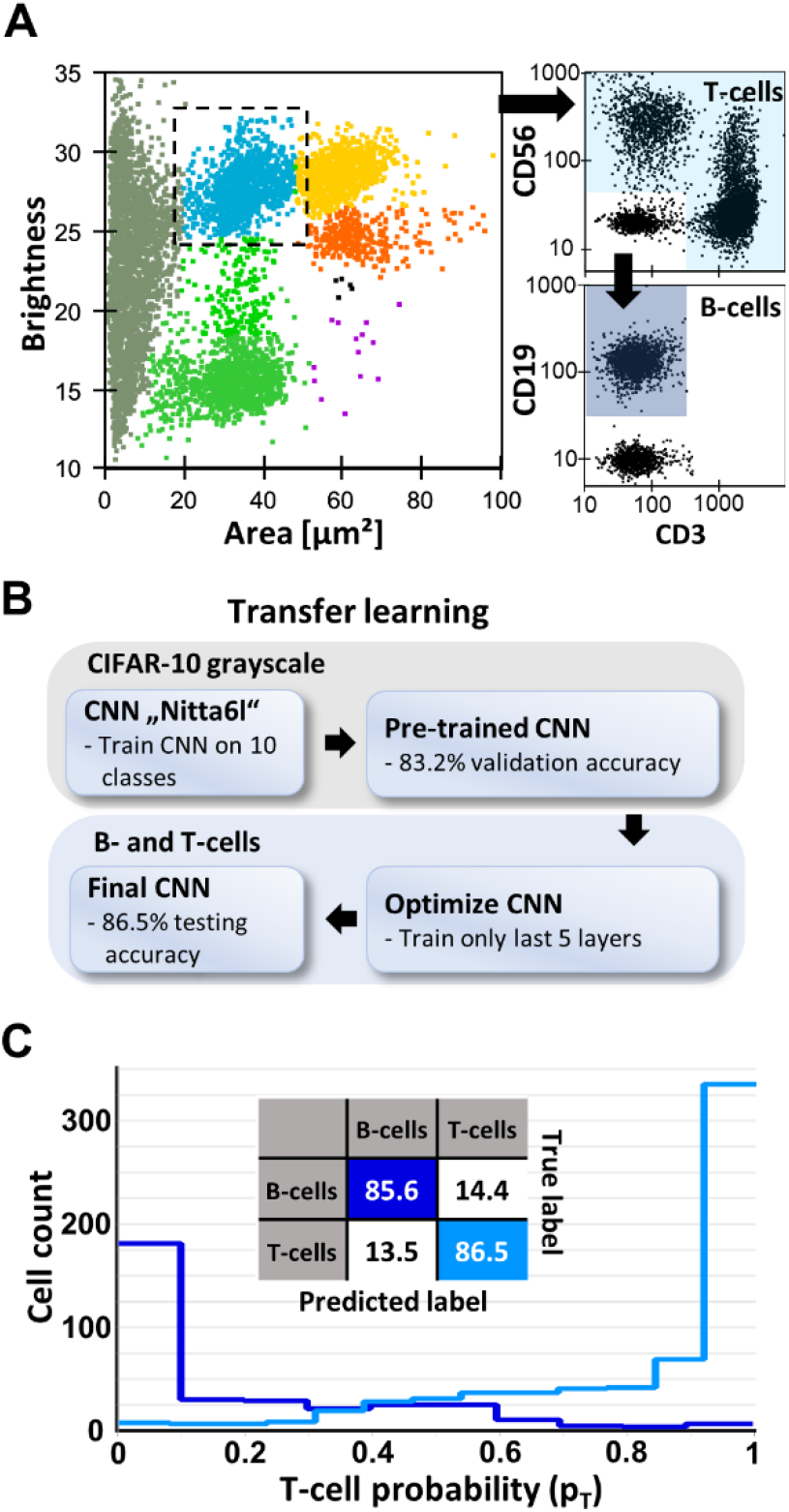
Label-free classification of B- and T-cells from human blood. (**A**) Gating strategy for acquiring training data for B- and T-cell classification. A scatterplot (brightness versus area) of human fractionated blood, measured using real-time deformability and fluorescence cytometry (RT-FDC) is shown. Lymphocytes were gated (dashed square) based on brightness and area (Toepfner et al., 2018). B- and T-cells were labeled according to standard surface CD markers (CD3 – T-cells, CD19 – B-cells and CD56 – NK-cells). (**B**) A representative schema of a transfer learning process, which can be easily applied in AID. The pretrained CNN_gray_, with a validation accuracy of 83.2% on the CIFAR-10 dataset, was loaded into AID and optimized to classify images of B- and T-cells, acquired from fractionated blood using RT-FDC. A final validation accuracy of 89.3% and a testing accuracy of 86.2% was achieved. (**C**) Confusion matrix of B-versus T-cells as well as the probability histogram showing the performance of the model on the testing set. The abscissa in the histogram shows the predicted probability to be a T-cell (p_T_).

## Discussion

### Graphical user interface provides simple access to deep learning methods

Advances in the quantity and quality of imaging data necessitate the availability and accessibility of processing tools, as automatization has the capacity to move scientific insights towards routine application. Numerous publications show the vast potential of DL for image processing, but ready-to-use software is not yet available (Shoham et al., 2018). Moreover, training of NNs requires expert programming skills thus limiting its accessibility for non-expert users. The open source community is capable to drive such software development, but the continuous evolution of programming environments and the fast turnover of software libraries impedes a cooperative progress. Furthermore, for inexperienced users the complex terminology regarding DL can prevent correct data assembly, successive training, and application of NNs. Some companies worked on user-friendly software solutions. Zeiss provides along with their microscopy software an addition called “Zen Intellesis”. This program can perform image feature extraction using a fixed NN, but does not support training of an individual NN. Instead, the extracted features are used to train a random forest (Breiman, 1996) for image segmentation. NVIDIA^®^ provides a user-interface called “DIGITS™” to assemble datasets and train NNs. However, DIGITS™ offers only limited access to hyper-parameters, no image augmentation methods, and only a few NNs which require specific input image dimensions. To adjust a model to other image dimensions or numbers of classes, coding is required. Further graphical user interface (GUI) based tools that apply machine learning to images are ilastik (Sommer et al., 2011), Fiji (Schindelin et al., 2012), and ImageJ (Schneider et al., 2012), but offer only limited access to DL methods.

AID addresses these issues by providing a standalone executable, including an intuitive GUI, which allows even non-programmers to train, evaluate, and apply NNs to their image datasets. Moreover, AID includes different NN architectures with different levels of complexity ranging from very simple multilayer perceptrons to contemporary CNNs with many layers. All these networks can be extended, updated, or replaced easily. Larger NNs might be favorable to maximize the classification accuracy while smaller models allow for real-time applications. Facilitating AID, label-free cell sorting for neutrophils from human blood was already demonstrated (Nawaz et al., 2019).

AID guides the user through the workflow to develop classification models, quantify their performance, and apply them to new data. Established methods for image normalization and augmentation are integrated. Interactive visualization tools allow a seamless link between user-settings and their effect on image data or the training process. Automatic documentation of the training process keeps track of all user-settings and model performance progression during training. Standard methods for quantification of model performance are embedded. Execution of AID is identical on Windows 7 and 10 PCs, allowing for reproducibility, sharing of models, and continuation of training using different PC setups. During creation of the standalone executable, we focused on achieving compatibility to a broad range of PC systems. Therefore, the leveraged DL Python library (TensorFlow) (Abadi et al., 2016) is restricted to use the central processing unit (CPU), as implementations to support a graphical processing unit (GPU) would lead to a dependency of AID on particular hardware. We demonstrated that considerable achievements with NNs are feasible despite the limitation to CPU processing. However, as AID is open source it can also be run from script which allows specific GPU support. Thus, AID could empower people to use DL for image classification, with implications for a wide range of disciplines, from life science to app development.

### Introducing AID using CIFAR-10 and Fashion-MNIST

We introduced the features of AID using CIFAR-10, a dataset of images commonly used as a standard to compare and benchmark image classification methods. We converted the RGB images to grayscale in order to train a model that can be re-used (“transfer learning”) for grayscale images retrieved from an imaging flow cytometer (RT-DC). Aided by manual optimization of hyper-parameters during the training process (Figure 1C), we reached a testing accuracy of 81.2%. Despite using a fairly low complex NN with only 4 convolutional layers and only grayscale information, our model would reach place 58 at https://benchmarks.ai/cifar-10. For comparison, we trained the same architecture using the original RGB images, resulting in a testing accuracy of 84.7%, corresponding to place 50 on https://benchmarks.ai/cifar-10. Furthermore, to show how models can be re-used, we applied transfer learning to optimize CNN_gray_ in order to classify images of ten fashion items (Fashion-MNIST) (Xiao et al., 2017). We obtained 95.1% validation and 93.8% testing accuracy, which to our knowledge is the highest testing accuracy ever reported for Fashion-MNIST. It should be noted that we did not develop a new NN architecture, but used a published architecture (Nitta et al., 2018) which we trained for multiple classification tasks.

Overall, AID can be used to load published NN architectures and train new models to classify both RGB and grayscale images. AID can convert RGB into grayscale upon user request. Furthermore, when creating a new NN, AID adjusts the input layer of the model according to the channel dimensions and user-defined input image size. Here, image input size is adjusted by center-cropping or padding. Currently, only squared input images are supported, but in the future rectangular image support will be implemented. Moreover, an interactive confusion matrix is displayed when a model is applied to new data (e.g. testing data), allowing the user to visualize correctly or incorrectly classified images.

### AID for life science applications and clinical blood diagnostics

Biomedical research and clinical diagnostics often utilize microscopy imaging, resulting in an increasing demand for automated image classification. While large tech-companies such as Google (https://deepmind.com/about/health) and Microsoft (https://www.microsoft.com/en-us/research/project/medical-image-analysis/) have initiated dedicated projects that focus on medical imaging, smaller research groups or start-up companies cannot always afford specialized staff for data analysis. Furthermore, enabling experts in a respective discipline to independently perform image analysis is advantageous for accurate data interpretation. Here, we demonstrate three applications where AID can be used in biomedical research: 1) quantification of adipogenic differentiation of MSCs from bright-field microscopy; 2) label-free blood count; and 3) label-free B-versus T-cell discrimination from imaging flow cytometry (RT-DC).

#### Quantification of adipogenic differentiation of MSCs

MSCs are a major source of stem cells used in cell therapy (Galipeau and Sensébé, 2018). Their differentiation potential is assessed by measuring areas of adipogenic differentiation in an MSC layer, which involves cell staining, imaging, and classification of the differentiated areas. Currently, computer-based quantification is challenging because of non-uniform staining, differences in cell morphology, and staining artifacts (Figure 2B and Figure 2—figure supplement 1). Therefore, we choose to train a NN to perform the detection of differentiated areas. More complex NNs have the capacity to learn more image characteristics, but as they typically contain more parameters, they tend to overfit (Srivastava et al., 2014). This issue can be tackled by increasing the amount of data. Here, due to the limited availability of labelled images, we applied a transfer learning approach, which was shown to reduce the need of data (Tan et al., 2018). We reached a validation accuracy of 92.1% by first training the model on CIFAR-10, and continuing training using the image data from differentiated MSCs. The validation results from 8 images confirmed that the model robustly quantified the differentiated areas, despite different imaging artifacts (Figure 2D and Figure 2—figure supplement 1). Application of this model to a classical research quantification tasked revealed that the model returned sensible predictions for new data (Krüger et al., 2020). The computational time required for classification (inference time) was approximately 4.5 ms for a single tile (32 x 32 pixels) and 0.4 s for a complete image (320 x 320 pixels) on an Intel^®^ Core™ i7-6800K CPU (3.4 GHz), rendering this model applicable for high-throughput analysis. It must be noted that AID is currently optimized for image classification problems and is not intended for image segmentation. In this study, we trained a binary classifier to predict whether individual tiles (32 x 32 pixels) from the entire image (320 x 320 pixels = 100 tiles) contain differentiated regions, or not. A more common practice is to train a model to return a segmentation map of the same size as the input image, allowing for pixel precise predictions (Badrinarayanan et al., 2015; Isola et al., 2016; Ronneberger et al., 2015).

#### Label-free blood count

Whereas manual microscopy can capture approximately 1 image per second, imaging cytometers easily speed up this task by a factor of 1000, resulting in considerably larger datasets. Here, we highlight the ability of AID to train CNNs on large datasets, by employing an imaging flow cytometer, RT-DC, to capture 1.2 million bright-field images of blood cells (Figure 3A). Cell subpopulations were gated manually based on brightness and area (Figure 3B; according to Toepfner et al., 2018) prior to loading into AID and performing training. This gating strategy implies that blood cell classification could be performed using area and brightness alone, however, these gates are not suitable for an automated classification strategy. This is due to the fact that brightness levels are highly dependent on experimental settings, such as focal plane, and therefore the brightness vs. area gate needs to be adjusted for each measurement. Our final CNN-based classification, in contrast, was trained to be robust to such technical alterations. Several image augmentation methods were used during training, including changes in orientation (rotation) and brightness levels to ensure robustness of the model. The final model was applied to classify RT-DC data from 17 individual whole blood measurements. The resulting blood count was very similar to a conventional clinical blood count (Figure 3C) indicating that the model is not influenced by mislabeled cells. This is expected since DL algorithms were shown to be robust against labelling noise, especially when using large datasets (Rolnick et al., 2017). The inference time for a single image is approximately 1 ms, corresponding to a prediction rate of 1,000 cells/s. which matches the image acquisition rate of RT-DC. Thus, in principle the model can be applied for on-the-fly prediction. RT-DC in combination with DL could complement the conventional whole blood count as we were also able to distinguish single red blood cells (RBCs) and assemblies of RBCs (doublets). RBC aggregates could be used as a diagnostic marker since their appearance is correlated to infection through increased fibrinogen concentration (Brust et al., 2015; Kamath and Lip, 2003). Furthermore, RBC aggregation is linked to erythrocyte sedimentation rate (ESR), which is a widely used marker to diagnose inflammatory or pathophysiological conditions (Bochen et al., 2001).

#### Label-free discrimination of B- and T-cells

Finally, we were able to train a CNN to distinguish B- and T-cells without label in RBC-depleted blood. We used transfer learning by loading the previously trained CNN_gray_ into AID and continued training using RT-FDC images of B- and T-cells. While CNN_gray_ was trained on a very different image dataset, elementary image features such as edges or corners also exist. Such simple image features are described by the first convolutional layer (Zeiler and Fergus, 2013) and we decided to omit this layer from training. AID offers this option within its user-interface. This strategy reduces the risk of overfitting and lowers the computational time since less parameters need to be updated during training. We continued training over the course of one month to promote a model with highest validation accuracy possible, demonstrating stable execution of AID for long run-times. Our results show that the classification performance of AID is at least similar to other publications showing labfel-free discrimination of B- and T-cells (Nassar et al., 2019; Yoon et al., 2018, 2017). These results suggest feasibility of label-free image-based discrimination of B- and T-cells, which could again be used to complement blood cell counts. Conventional methods for classifying these cells involve fluorescent labelling and sorting which are time-consuming and costly. Moreover, molecular labelling involves the risk of contamination, limiting the use of the cells after sorting. Given the large margin between the predicted probabilities of B- and T-cells (Figure 4C), combining label-free image-based sorting and this model could result in highly pure B- or T-cells samples. Promising technologies to perform this task were introduced recently (Nawaz et al., 2019; Nitta et al., 2018).

In conclusion, we present a software tool which drastically increases the accessibility of AI-based image classification to non-experts. AID can handle RGB and grayscale images in small and large datasets and has state-of-the-art techniques for improved training of NNs, such as image augmentation and transfer learning already implemented. We have demonstrated the power of AID for obtaining robust classifiers for multiple use-cases covering a wide spectrum of applications. This shows that AID is ready to be applied by anyone to image classification problems in life science and beyond.

## Materials und Methods

### Computer equipment and software

AIDeveloper (AID) is open source (https://github.com/maikherbig/AIDeveloper). AID was written in Python 3.5 using PyQT (package for graphical user interface), Keras (https://keras.io/) and TensorFlow (Abadi et al., 2016) (packages for deep learning) and several further open source Python packages (Supplementary file 1). We used PyInstaller to freeze AID to a standalone executable, running on Windows 7 and 10 PCs. While TensorFlow can be installed to support graphical processing unit (GPU) usage, such an installation would then depend on particular GPU hardware, limiting the compatibility of AID with other PC systems. Hence, we restricted the frozen AID software to use CPU implementations of TensorFlow, which was exclusively used throughout this work. For training neural nets, we used a PC system with an Intel^®^ Core™ i7-3930K CPU @ 3.2 GHz and 32 GB RAM.

### CIFAR-10

CIFAR-10 is a labelled dataset containing 60,000 RGB-images of 10 different classes (Krizhevsky, 2009). We downloaded the dataset from https://pjreddie.com/projects/cifar-10-dataset-mirror/. In this dataset, 50,000 images are dedicated to training and 10,000 to testing. We picked 200 random training images of each class to create a validation set. To convert images to grayscale, AID uses the luminosity method, which performs a weighted average of the channels of an RGB image, accounting for the higher sensitivity of the human eye to green color: gray=0.21×R+0.72×G×0.07×B.

### Mesenchymal stromal cell isolation, culture, differentiation and dataset acquisition

MSC isolation was performed in compliance with the Declaration of Helsinki. Bone marrow (BM) aspirates were taken from healthy volunteer donors, after obtaining informed written consent, and cells were isolated (ethical approval no. EK221102004, EK47022007) according to a previously reported method (Majumdar et al., 1998). In brief, BM aspirates were diluted in phosphate buffered saline (PBS, Sigma-Aldrich, Germany) at a ratio of 1:2 and a density gradient centrifugation (2,000 g for 15 min at room temperature) using a 20 mL aliquot layered over a 1.073 g/mL Percoll solution (Biochrom, Berlin, Germany). The fraction of mononucleated cells (MNCs) were transferred to a cell culture flask in MSC medium. MSC medium was prepared using Dulbecco’s modified Eagle medium (DMEM) with glucose and L-glutamine (BioWhittaker^®^ DMEM Culture Media, VWR, Dresden, Germany) and 10% foetal calf serum (FCS). MSCs were detached using trypsin and transferred to 3 wells of a 6-well culture plate. Adipogenesis differentiation was induced when cells reached 80% confluency as previously described (Oswald et al., 2004). Briefly, adipogenesis was induced with 1 μmol/L dexamethasone, 0.5 mmol/L 3-isobutyl-1-methylxanthine, 100 μmol/L indomethacin, 10 μmol/L insulin (Sigma-Aldrich, St. Louis, USA) and 10% FCS in DMEM for 14 days. All cultures were kept at 37 °C with 5% CO_2_ in a water-jacked incubator. Medium changes were performed weekly. For histological visualization differentiated cells were fixed with 4 % paraformaldehyde (Merck KGaA, Darmstadt, Germany) in PBS. Adipogenic differentiation was assessed by Oil Red O staining with 0.1% Oil Red O solution (Sigma-Aldrich, St. Louis, USA), followed by five washes with distilled water. To generate the image dataset an inverted microscope (Axiovert 25, Carl-Zeiss, Jena, Germany) equipped with a digital camera (Olympus E330, Olympus, Hamburg, Germany) was used. Images from 5 different positions of each well were taken (Figure 3A, B).

Labeling was performed by an expert marking each pixel that corresponded to an area of differentiated cells, resulting in a pixel precise map. In total, 46 images from 16 different donors were labelled. 38 labeled images were used for training and the remaining 8 images for validation. Each original image of 320 x 320 pixels in size was partitioned into 100 tiles of 32 x 32 pixels. The chosen size of 32 x 32 pixels approximately meets the size of the objects and also reflects a compromise between large tiles which would often contain several objects and single-pixel-tiles that would prevent a model from learning about the morphology of the objects. A tile was assigned to class 0 (“without differentiation”), if it contained less than five marked pixels or to class 1 (“with differentiation”) if it contained more or equal to 5 pixels. After equally partitioning all images, tiles of class 1 were clearly under-represented. Therefore, more tiles from random locations containing more than 4 marked pixels were added. This strategy helps to balance the dataset and allows to obtain tiles with objects at various locations in the image. The latter can help to train a more translation invariant model.

### Real-time fluorescence and deformability cytometry for blood

Real-time deformability and fluorescence cytometry was performed as described elsewhere (Otto et al., 2015; Rosendahl et al., 2018). Briefly, a microfluidic chip made from polydimethylsiloxane (PDMS, SYLGARD^®^, Dow Corning, USA) was mounted on an inverted microscope (Observer Z1, Zeiss, Jena, Germany) equipped with an LED (CBT-120, Luminus Devices, USA) and a high-speed camera (EoSens CL MC1362, Mikrotron, Unterschleißheim, Germany). Two syringe pumps (NemeSyS, Cetoni, Germany) were used to deliver suspended cells and sheath fluid into the chip at a total flow rate of 0.06 μL/s. Cells were captured inside a constriction channel of 20 μm x 20 μm cross-section (Figure 4A). For suspending cells, a measurement buffer (MB) based on Mg^2+^-and Ca^2+^-free PBS, supplemented with 0.6% (w/w) methylcellulose, was used.

For preparation of whole blood samples, venous blood was drawn from human donors using a 20-gauge multifly needle into sodium citrate tubes (S-Monovette^®^ 10 mL 9NC, Sarstedt, Germany) by vacuum aspiration. Whole blood samples were prepared by diluting 50 μL of whole blood in 950 μL of MB as previously published (Toepfner et al., 2018).

To prepare RBC-depleted blood samples, 2 mL of a 6% dextran solution (Dextran T500, Pharmacosmos A/S, Denmark) diluted in sodium chloride (0.9% Sodium Chloride Irrigation, Baxter Healthcare, Switzerland) was added to 10 mL of whole citrated blood. After gentle mixing, RBCs were allowed to sediment for 30 min (Bøyum, 2008; Quach and Ferrante, 2017). The supernatant was transferred to a 10 mL tube and centrifuged for 10 min at 120 g (Universal 30RF, Hettich, Switzerland). After removing the cell-free plasma, the pellet was resuspended in 2 mL MB.

For B- and T-cell classification fractionated blood was used. Blood aspirates were diluted in PBS at a ratio of 1:5, followed by a density gradient centrifugation (2000 g for 15 min at room temperature) using a 20 mL aliquot layered over a 1,073 g/mL Percoll solution (Biochrom, Berlin, Germany). The MNC fraction was washed once in PBS and 100 μL aliquots were used to stain B-cells using an antibody against CD19 (coupled to allophycocyanin; APC), T-cells using anti-CD3 (coupled to fluorescein; FITC) and NK-cells, as a subset of T-cells using anti-CD56 (coupled to phycoerythrin; PE). Cells were washed in PBS and finally resuspended in MB. Real-time fluorescence and deformability cytometry (RT-FDC) was performed as described elsewhere (Rosendahl et al., 2018). Briefly, cells were flushed through a constriction in a microfluidic chip at a flowrate if 0.06 μL/s. A laser sheet was projected into the middle of the channel. When cells passed through the sheet, three lasers (488nm, 561nm, 640nm) excited the fluorescence signal and the fluorescence intensity was measured by dedicated detectors; resulting in three 1D-fluorescence traces for each cell. Here, we used the maximum peak-height of the fluorescence traces to quantify whether the corresponding cell is expressing a particular fluorescent marker. Since bright-field image and fluorescence acquisition are synchronized, the fluorescence information can be used as ground truth to label each image.

All studies complied with the Declaration of Helsinki and involved written informed consent from all participants. Donors were recruited at the University Medical Centre Carl Gustav Carus Dresden and ethics for experiments with human blood were approved by the ethics committee of the Technische Universität Dresden (EK89032013, EK458102015).

## Acknowledgement

We thankfully acknowledge the multitude of feedback and ideas for improvement of the practical user-experience of AID from Karen Teßmer and Felix Reichel. Furthermore, we thankfully received support and advice regarding icons and design elements of AID from Konrad Wauer. The authors acknowledge funding from the Alexander von Humboldt-Stiftung (Alexander von Humboldt Professorship to J.G.), Deutsche Forschungsgesellschaft (DFG Project number 399422891 to J.G. and Marius Ader), Marie Sklodowska-Curie Actions under the European Union’s Horizon 2020 research and innovation programme (grant number 641639 to S.A. and J.G.) and a DKMS ‘Mechthild Harf Research Grant’ (DKMS-SLS-MHG-2016-02 to A.J.).

## Author contributions

M.H. and M.K. conceived the project, elected the experiments and conducted training of neural nets in AID. M.H. developed the software AIDeveloper. M.H, M.K., A.J. and T.K performed experiments. M.K. conceived and performed the RT-FDC experiment for B and T-cells discrimination as well as the corresponding CNN training in AID. M.K. visualized the data and prepared all figures. M.H., M.K., S.A. and D.S. co-wrote the manuscript. All authors reviewed the manuscript.

## Supplementary Figures

**Figure 1—figure supplement 1:**
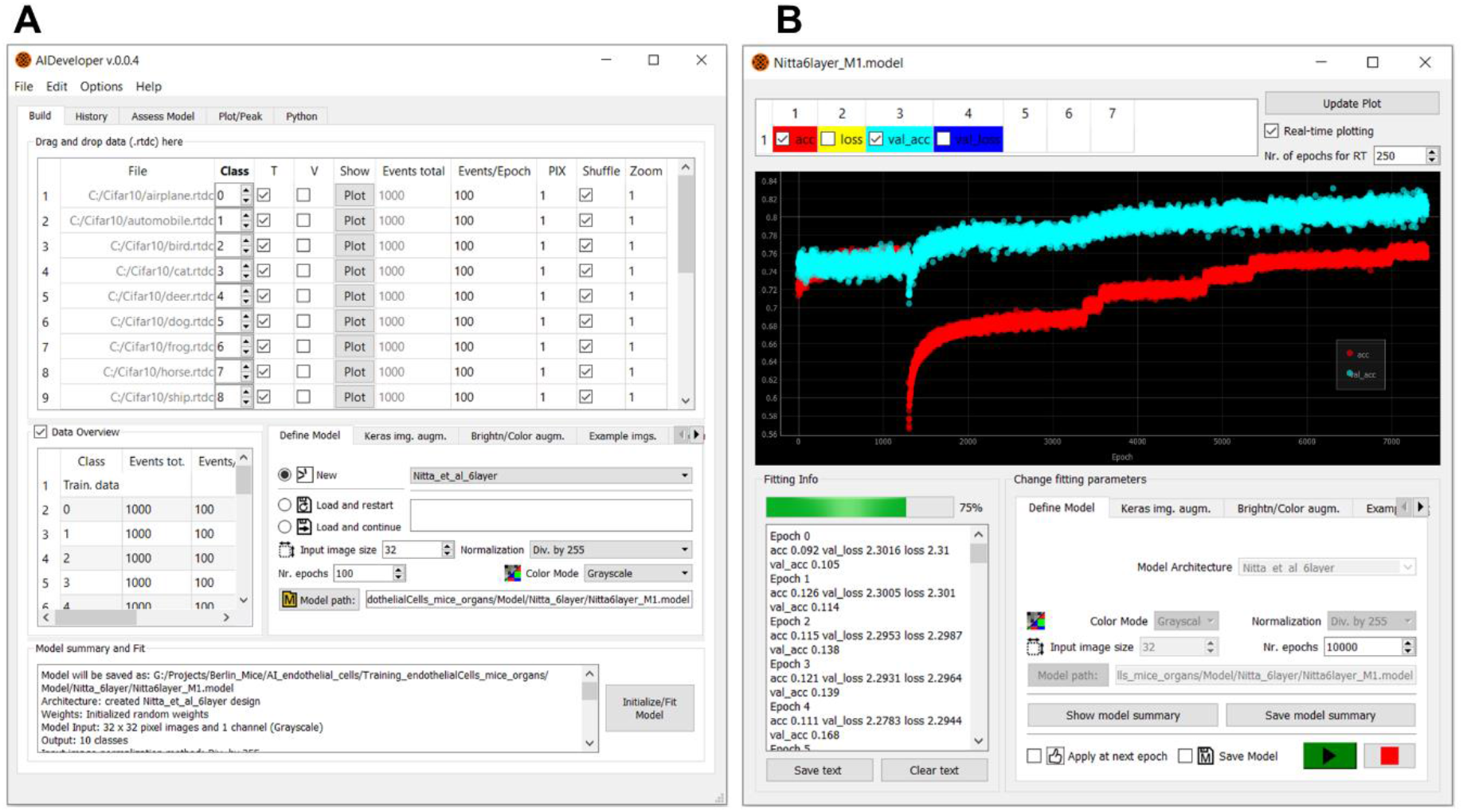
Screenshots of the graphical user interface (GUI) of AID. Intuitive elements allow the user to load data, select a neural net, and set hyper-parameters. (**A**) Main user interface of AID which shows a table of the loaded data and allows the user to define classes and which dataset belongs to training and validation set. Definition of the model is performed by choosing an architecture and input image size in corresponding GUI elements. (**B**) Real-time visualization of the progress during training of a specific model showing accuracy (red) and validation accuracy (cyan).

**Figure 1—figure supplement 2:**
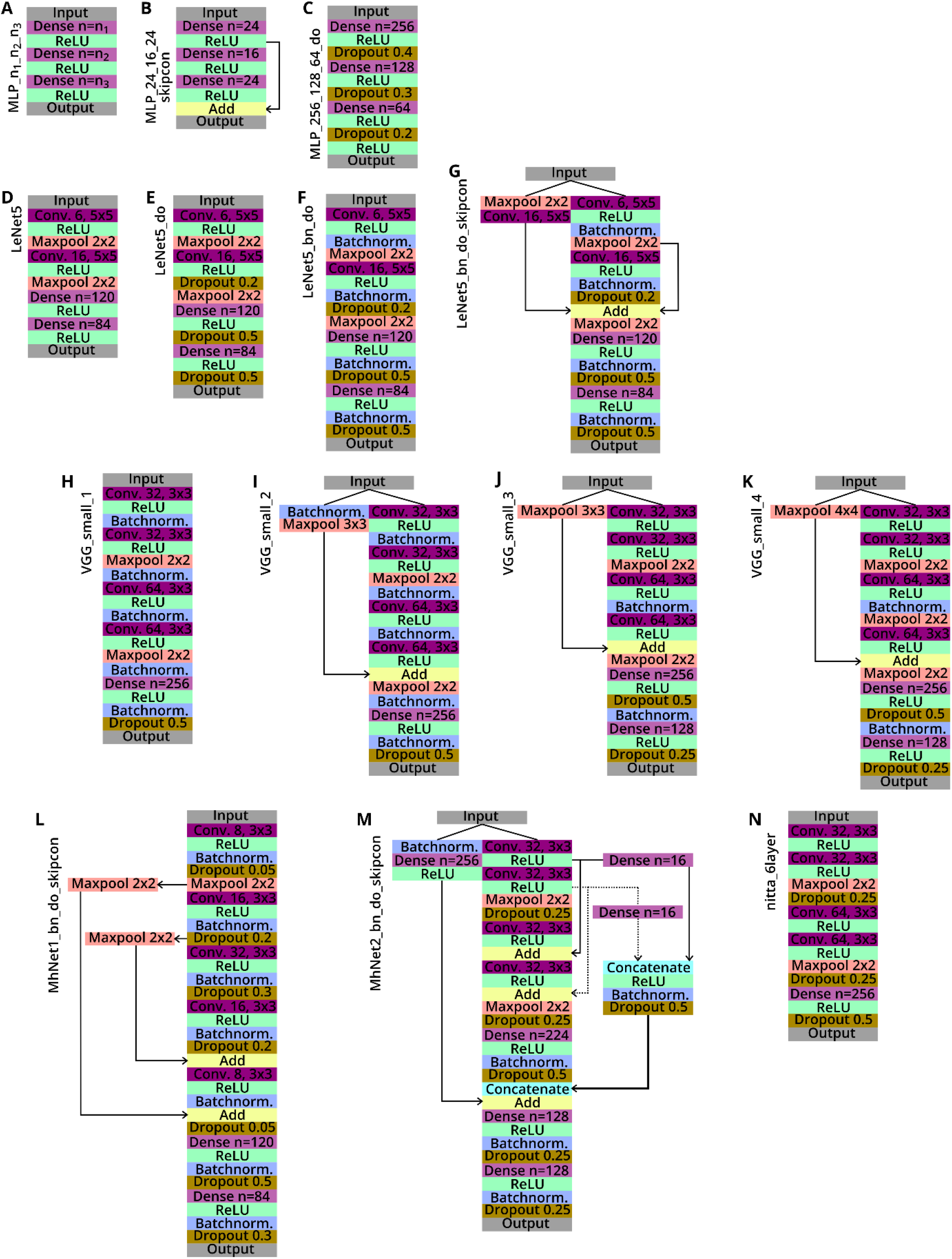
Sketches of neural nets (NN) that are implemented in AIDeveloper. Each coloured box represents an individual layer of the network. Input layers in AID currently only accept squared images with either one (grayscale) or three channels (RGB image). The output layer always consists of a fully connected layer including as many nodes as different output classes and a softmax activation layer (Goodfellow et al., 2016, p. 184). Fully connected layers are abbreviated ‘Dense’ and the number of nodes is given. In all shown NNs, rectified linear units (ReLU) are used as activation function. Convolutional layers are abbreviated ‘Conv.’ and the number of convolutional filters and the filter-size is given. Dropout layers are shown including their respective dropout rate. Downsampling is achieved using maxpooling layers and the pooling size is shown. Arrows indicate skip-connections, which bypass the main thread of the NN and either add or concatenate their data back to the NN in a subsequent layer.

**Figure 1—figure supplement 3:**
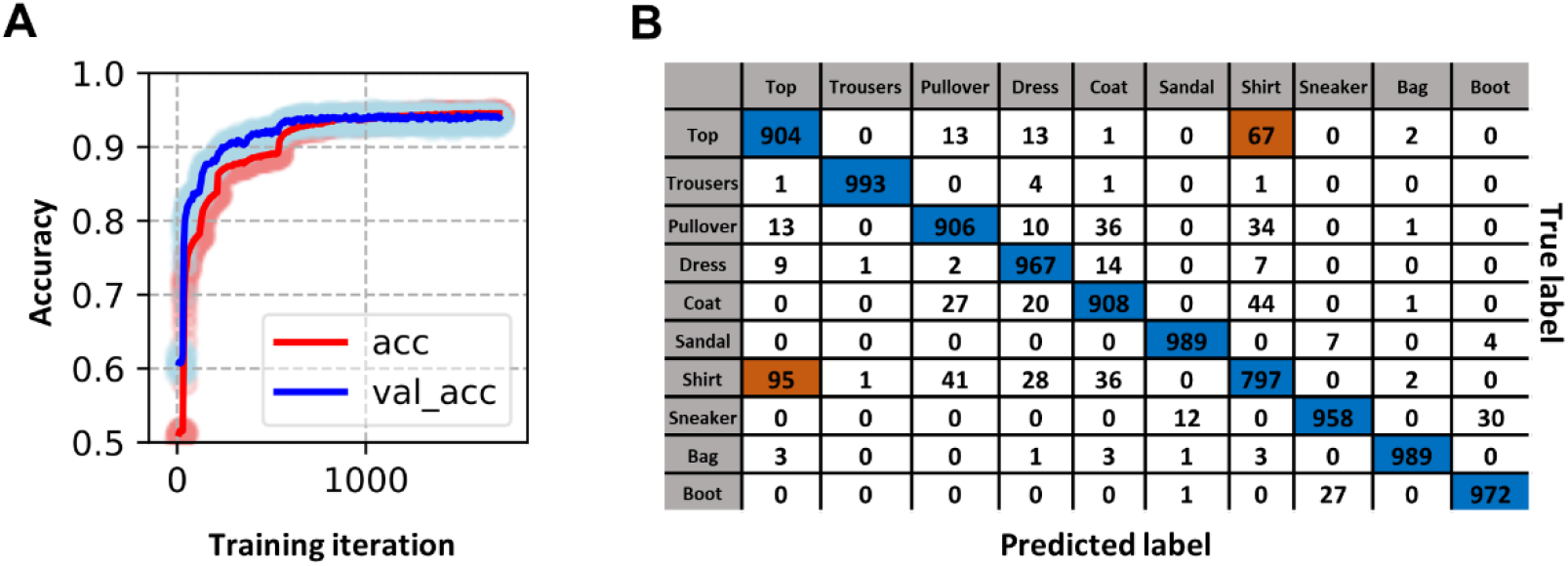
Transfer learning approach for Fashion-MNIST. (**A**) The image shows a training process for the classification of fashion items from the Fashion-MNIST dataset. Here, transfer learning was applied by re-using a model (CNN_gray_) previously trained on CIFAR-10. The red line (acc) shows the rolling median of the accuracy (window size = 10 training iterations) and light red indicates the accuracy of individual epochs. The blue line (val_acc) shows the corresponding rolling median of the validation accuracy and light blue indicates the validation accuracy of individual epochs. The model with the highest validation accuracy (95.1%) was applied to the testing set resulting in a testing accuracy of 93.8%. (**B**) The confusion matrix shows the performance of the final model on the testing set. The model appears to have most difficulties in distinguishing “Top” and “Shirt” (highlighted in orange).

**Figure 2—figure supplement 3:**
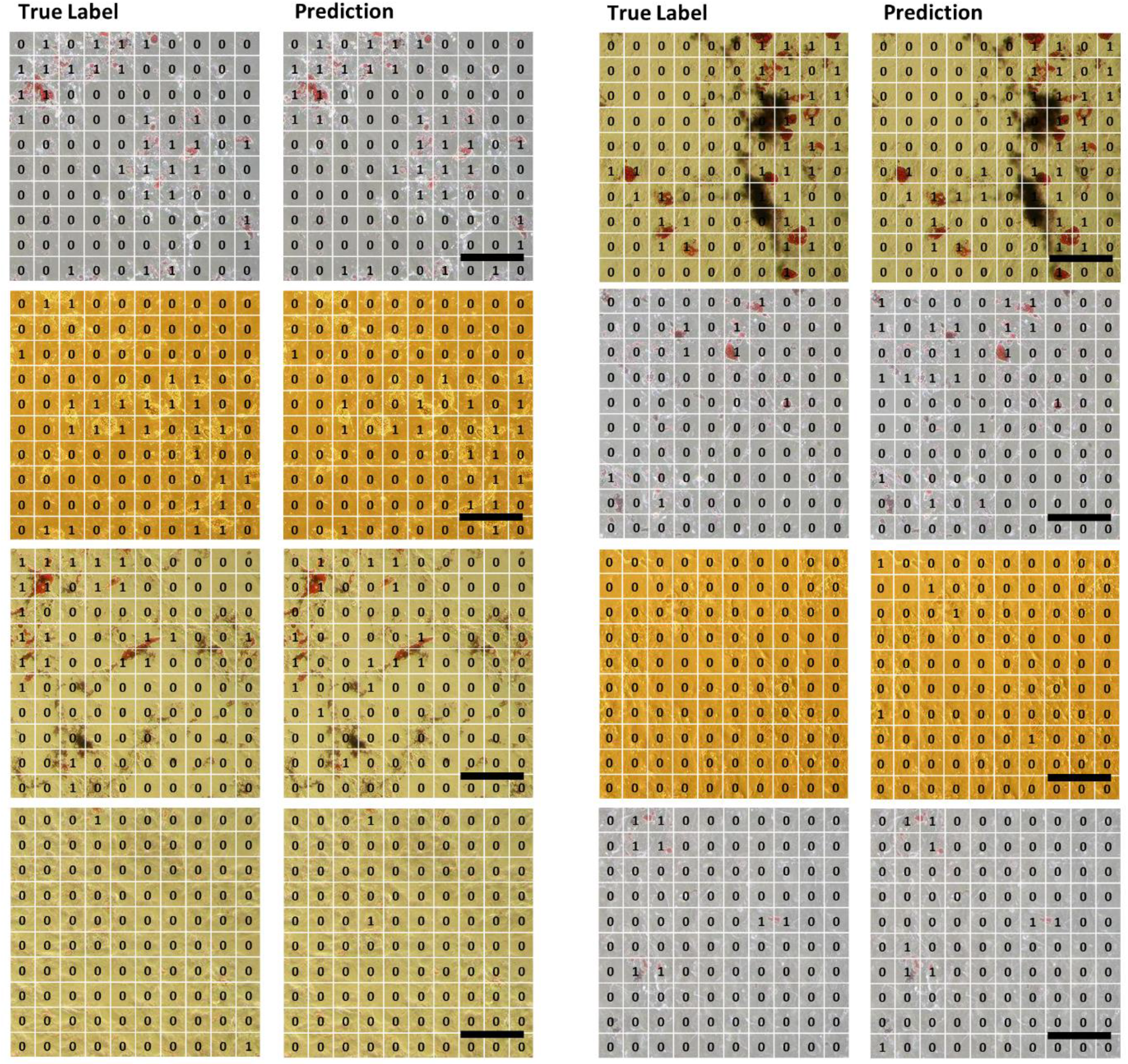
Adipocyte detection in validation images. The micrographs of MSC layers induced to differentiate into the adipogenic direction highlight the image colour and Oil Red O staining variability. Regions in the image containing differentiated adipocytes were marked by an expert (True label, left column). The trained neural net was applied to predict those regions (Predicted label, right column). Scale bars = 50 μm.

## Supplementary file 1: Python libraries used

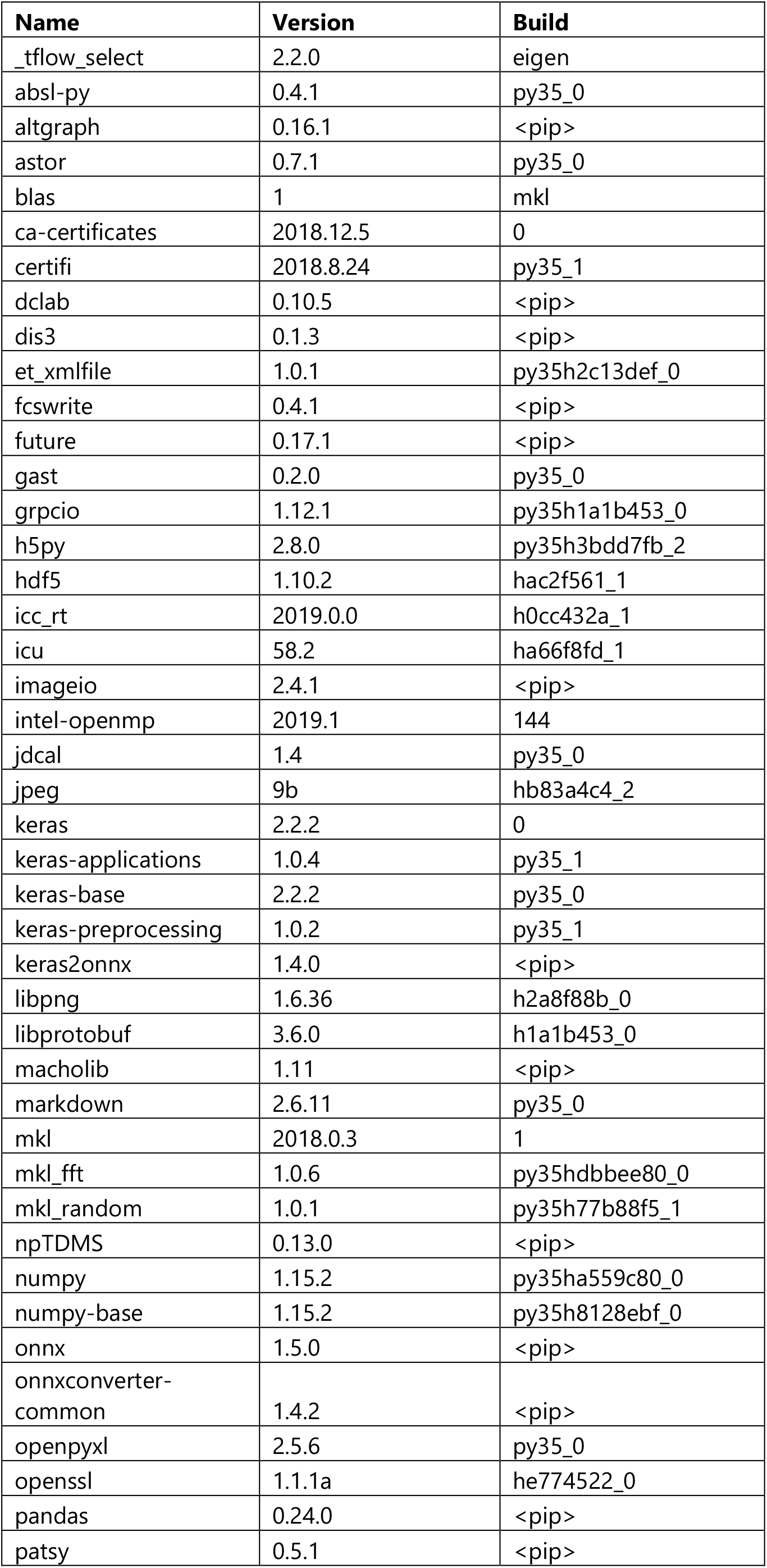
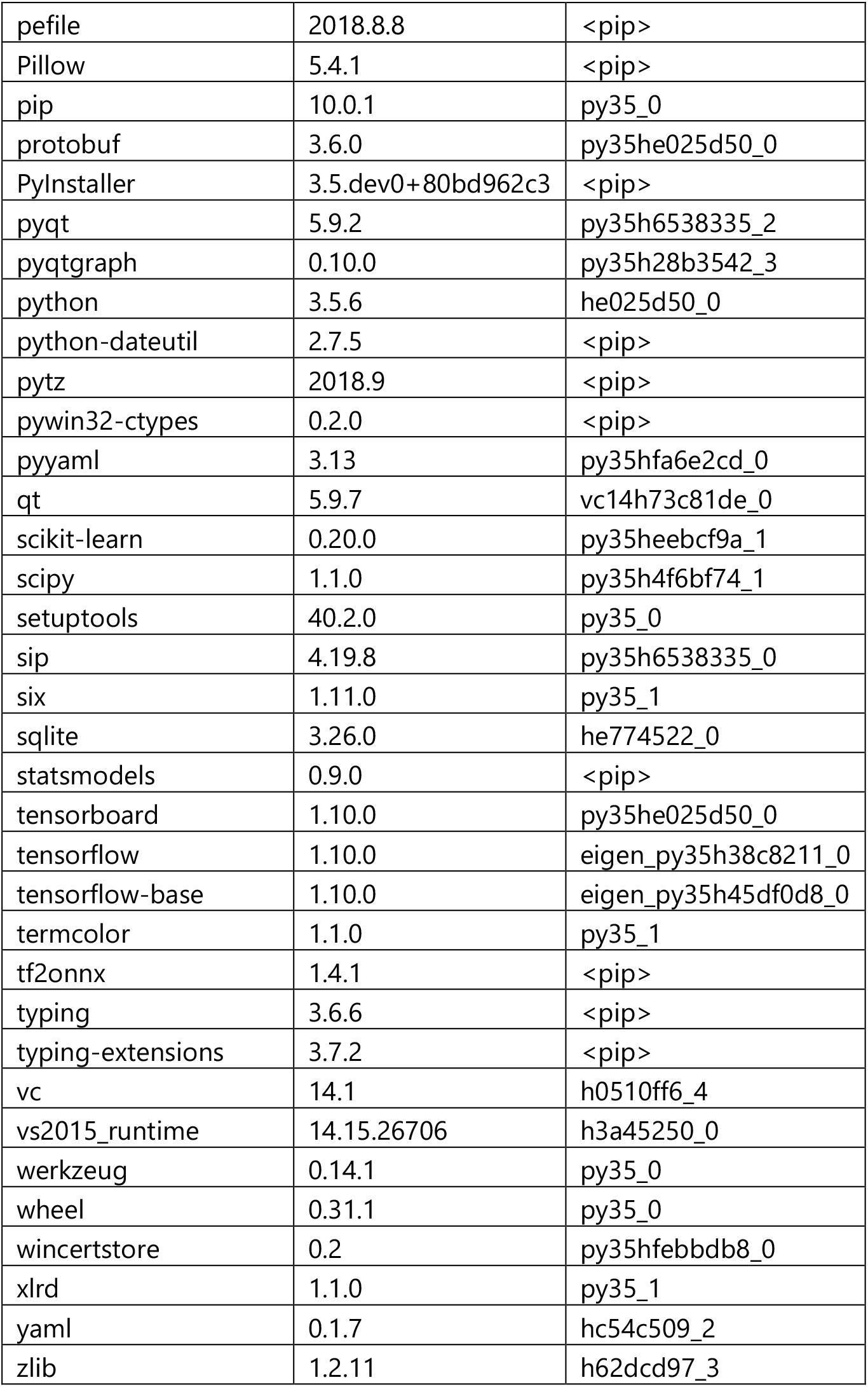

## References

Abadi M, Agarwal A, Barham P, Brevdo E, Chen Z, Citro C, Corrado GS, Davis A, Dean J, Devin M, Ghemawat S, Goodfellow I, Harp A, Irving G, Isard M, Jia Y, Jozefowicz R, Kaiser L, Kudlur M, Levenberg J, Mane D, Monga R, Moore S, Murray D, Olah C, Schuster M, Shlens J, Steiner B, Sutskever I, Talwar K, Tucker P, Vanhoucke V, Vasudevan V, Viegas F, Vinyals O, Warden P, Wattenberg M, Wicke M, Yu Y, Zheng X. 2016. TensorFlow: Large-Scale Machine Learning on Heterogeneous Distributed Systems.

Badrinarayanan V, Kendall A, Cipolla R. 2015. SegNet: A Deep Convolutional Encoder-Decoder Architecture for Image Segmentation.

Bochen K, Krasowska A, Milaniuk S, Kulczyńska M, Prystupa A, Dzida G. 2001. Erythrocyte sedimentation rate – an old marker with new applications. J Pre-Clinical Clin Res 5:50–55.

Bøyum A. 2008. Separation of Blood Leucocytes, Granulocytes and Lymphocytes. Tissue Antigens 4:269–274. doi:10.1111/j.1399-0039.1974.tb00252.x

Breiman L. 1996. Bagging Predictors. Mach Learn 24:123–140. doi:10.1023/A:1018054314350

Brust M, Aouane O, Thiébaud M, Flormann D, Verdier C, Kaestner L, Laschke MW, Selmi H, Benyoussef A, Podgorski T, Coupier G, Misbah C, Wagner C. 2015. The plasma protein fibrinogen stabilizes clusters of red blood cells in microcapillary flows. Sci Rep 4:4348. doi:10.1038/srep04348

Buggenthin F, Buettner F, Hoppe PS, Endele M, Kroiss M, Strasser M, Schwarzfischer M, Loeffler D, Kokkaliaris KD, Hilsenbeck O, Schroeder T, Theis FJ, Marr C. 2017. Prospective identification of hematopoietic lineage choice by deep learning. Nat Methods 1–7. doi:10.1038/nmeth.4182

Caesar H, Bankiti V, Lang AH, Vora S, Liong VE, Xu Q, Krishnan A, Pan Y, Baldan G, Beijbom O. 2019. nuScenes: A multimodal dataset for autonomous driving.

Cireşan DC, Giusti A, Gambardella LM, Schmidhuber J. 2013. Mitosis Detection in Breast Cancer Histology Images with Deep Neural Networks. Springer, Berlin, Heidelberg. pp. 411–418. doi:10.1007/978-3-642-40763-5_51

Esteva A, Kuprel B, Novoa RA, Ko J, Swetter SM, Blau HM, Thrun S. 2017. Dermatologist-level classification of skin cancer with deep neural networks. Nature 542:115–118. doi:10.1038/nature21056

Galipeau J, Sensébé L. 2018. Mesenchymal Stromal Cells: Clinical Challenges and Therapeutic Opportunities. Cell Stem Cell 22:824–833. doi:10.1016/j.stem.2018.05.004

Giusti A, Caccia C, Ciresari DC, Schmidhuber J, Gambardella LM. 2014. A comparison of algorithms and humans for mitosis detection2014 IEEE 11th International Symposium on Biomedical Imaging (ISBI). IEEE. pp. 1360–1363. doi:10.1109/ISBI.2014.6868130

Goodfellow I, Bengio Y, Courville A. 2016. Deep learning. MIT Press.

Isola P, Zhu J-Y, Zhou T, Efros AA. 2016. Image-to-Image Translation with Conditional Adversarial Networks.

Jones TR, Kang I, Wheeler DB, Lindquist RA, Papallo A, Sabatini DM, Golland P, Carpenter AE. 2008. CellProfiler Analyst: data exploration and analysis software for complex image-based screens. BMC Bioinformatics 9:482. doi:10.1186/1471-2105-9-482

Kamath S, Lip GYH. 2003. Fibrinogen: biochemistry, epidemiology and determinants. QJM 96:711–729. doi:10.1093/qjmed/hcg129

Krizhevsky A. 2009. Learning Multiple Layers of Features from Tiny Images.

Krüger T, Middeke JM, Stölzel F, Mütherig A, List C, Brandt K, Heidrich K, Teipel R, Ordemann R, Schuler U, Oelschlägel U, Wermke M, Kräter M, Herbig M, Wehner R, Schmitz M, Bornhäuser M, von Bonin M. 2020. Reliable isolation of human mesenchymal stromal cells from bone marrow biopsy specimens in patients after allogeneic hematopoietic cell transplantation. Cytotherapy 22:21–26. doi:10.1016/j.jcyt.2019.10.012

Lecun Y, Bottou L, Bengio Y, Haffner P. 1998. Gradient-based learning applied to document recognition. Proc IEEE 86:2278–2324. doi:10.1109/5.726791

Majumdar MK, Thiede MA, Mosca JD, Moorman M, Gerson SL. 1998. Phenotypic and functional comparison of cultures of marrow-derived mesenchymal stem cells (MSCs) and stromal cells. J Cell Physiol 176:57–66. doi:10.1002/(SICI)1097-4652(199807)176:1<57::AID-JCP7>3.0.CO;2-7

Nassar M, Doan M, Filby A, Wolkenhauer O, Fogg DK, Piasecka J, Thornton CA, Carpenter AE, Summers HD, Rees P, Hennig H. 2019. Label-Free Identification of White Blood Cells Using Machine Learning. Cytom Part A cyto.a. 23794. doi:10.1002/cyto.a.23794

Nawaz AA, Urbanska M, Herbig M, Nötzel M, Kräter M, Rosendahl P, Herold C, Toepfner N, Kubankova M, Goswami R, Abuhattum S, Reichel F, Müller P, Taubenberger A, Girardo S, Jacobi A, Guck J. 2019. Using real-time fluorescence and deformability cytometry and deep learning to transfer molecular specificity to label-free sorting. bioRxiv 862227. doi:10.1101/862227

Nitta N, Sugimura T, Isozaki A, Mikami H, Hiraki K, Sakuma S, Iino T, Arai F, Endo T, Fujiwaki Y, Fukuzawa H, Hase M, Hayakawa T, Hiramatsu K, Hoshino Y, Inaba M, Ito T, Karakawa H, Kasai Y, Koizumi K, Lee SW, Lei C, Li M, Maeno T, Matsusaka S, Murakami D, Nakagawa A, Oguchi Y, Oikawa M, Ota T, Shiba K, Shintaku H, Shirasaki Y, Suga K, Suzuki Y, Suzuki N, Tanaka Y, Tezuka H, Toyokawa C, Yalikun Y, Yamada M, Yamagishi M, Yamano T, Yasumoto A, Yatomi Y, Yazawa M, Di Carlo D, Hosokawa Y, Uemura S, Ozeki Y, Goda K. 2018. Intelligent Image-Activated Cell Sorting. Cell 175:266–276.e13. doi:10.1016/j.cell.2018.08.028

Oswald J, Boxberger S, Jørgensen B, Feldmann S, Ehninger G, Bornhäuser M, Werner C. 2004. Mesenchymal Stem Cells Can Be Differentiated Into Endothelial Cells In Vitro. Stem Cells 22:377–384. doi:10.1634/stemcells.22-3-377

Otto O, Rosendahl P, Mietke A, Golfier S, Herold C, Klaue D, Girardo S, Pagliara S, Ekpenyong A, Jacobi A, Wobus M, Töpfner N, Keyser UF, Mansfeld J, Fischer-Friedrich E, Guck J. 2015. Real-time deformability cytometry: on-the-fly cell mechanical phenotyping. Nat Methods 12. doi:10.1038/nmeth.3281

Quach A, Ferrante A. 2017. The Application of Dextran Sedimentation as an Initial Step in Neutrophil Purification Promotes Their Stimulation, due to the Presence of Monocytes. J Immunol Res 2017:1–10. doi:10.1155/2017/1254792

Rolnick D, Veit A, Belongie S, Shavit N. 2017. Deep Learning is Robust to Massive Label Noise.

Ronneberger O, Fischer P, Brox T. 2015. U-Net: Convolutional Networks for Biomedical Image Segmentation.

Rosendahl P, Plak K, Jacobi A, Kraeter M, Toepfner N, Otto O, Herold C, Winzi M, Herbig M, Ge Y, Girardo S, Wagner K, Baum B, Guck J. 2018. Real-time fluorescence and deformability cytometry. Nat Methods. doi:10.1038/nmeth.4639

Schindelin J, Arganda-Carreras I, Frise E, Kaynig V, Longair M, Pietzsch T, Preibisch S, Rueden C, Saalfeld S, Schmid B, Tinevez J-Y, White DJ, Hartenstein V, Eliceiri K, Tomancak P, Cardona A. 2012. Fiji: an open-source platform for biological-image analysis. Nat Methods 9:676–682. doi:10.1038/nmeth.2019

Schneider CA, Rasband WS, Eliceiri KW. 2012. NIH Image to ImageJ: 25 years of image analysis. Nat Methods 9:671–675. doi:10.1038/nmeth.2089

Shoham Y, Perrault R, Brynjolfsson E, Clark J, Manyika J, Niebles JC, Lyons T, Etchemendy J, Grosz B, Bauer Z. 2018. The AI Index 2018 Annual Report. Stanford, CA.

Sommer C, Straehle C, Kothe U, Hamprecht FA. 2011. Ilastik: Interactive learning and segmentation toolkit2011 IEEE International Symposium on Biomedical Imaging: From Nano to Macro. IEEE. pp. 230–233. doi:10.1109/ISBI.2011.5872394

Srivastava N, Hinton G, Krizhevsky A, Sutskever I, Salakhutdinov R. 2014. Dropout: A Simple Way to Prevent Neural Networks from Overfitting. J Mach Learn Res 15:1929–1958.

Swets JA. 1996. Signal detection theory and ROC analysis in psychology and diagnostics: collected papers. L. Erlbaum Associates.

Tan C, Sun F, Kong T, Zhang W, Yang C, Liu C. 2018. A Survey on Deep Transfer Learning.

Toepfner N, Herold C, Otto O, Rosendahl P, Jacobi A, Kräter M, Stächele J, Menschner L, Herbig M, Ciuffreda L, Ranford-Cartwright L, Grzybek M, Coskun Ü, Reithuber E, Garriss G, Mellroth P, Henriques-Normark B, Tregay N, Suttorp M, Bornhäuser M, Chilvers ER, Berner R, Guck J. 2018. Detection of human disease conditions by single-cell morpho-rheological phenotyping of blood. Elife 7:1–22. doi:10.7554/eLife.29213

Xiao H, Rasul K, Vollgraf R. 2017. Fashion-MNIST: a Novel Image Dataset for Benchmarking Machine Learning Algorithms.

Yoon J, Jo Y, Kim M, Kim K, Lee S, Kang S-J, Park Y. 2017. Identification of non-activated lymphocytes using three-dimensional refractive index tomography and machine learning. Sci Rep 7:6654. doi:10.1038/s41598-017-06311-y

Yoon J, Jo Y, Kim YS, Yu Y, Park J, Lee S, Park WS, Park Y. 2018. Label-Free Identification of Lymphocyte Subtypes Using Three-Dimensional Quantitative Phase Imaging and Machine Learning. J Vis Exp. doi:10.3791/58305

Yosinski J, Clune J, Bengio Y, Lipson H. 2014. How transferable are features in deep neural networks?

Zeiler MD, Fergus R. 2013. Visualizing and Understanding Convolutional Networks.

